# Aberrant structural synaptic dynamics precede disorganization of temporal and spatial coding in the hippocampus upon repeated stress exposure

**DOI:** 10.1101/2020.11.30.403972

**Authors:** Alireza Chenani, Ghabiba Weston, Alessandro F. Ulivi, Tim P. Castello-Waldow, Alon Chen, Alessio Attardo

**Author notes:** Equal contribution (Alireza Chenani, Ghabiba Weston).

## Abstract

Stress exposure strongly contributes to the pathophysiology of psychiatric diseases such as depression, schizophrenia, anxiety, and post-traumatic stress disorder. It also affects both function and structure of hippocampal CA1, leading to episodic memory impairment. Here, we used deep-brain optical imaging to elucidate the effects of stress on CA1 pyramidal neuron structural connectivity and activity.

We tracked dynamics of dendritic spines during repeated stress and found decrease in spinogenesis followed by decrease in spine stability. In contrast to repeated stress, acute stress led to stabilization of the spines born in temporal proximity to the stressful event. To investigate the link between structural plasticity and activity patterns upon repeated stress, we studied the activity of thousands of CA1 pyramidal neurons in freely-moving. We found an increase in activity and loss of temporal organization followed by a disruption in temporal and spatial coding. Our data suggest that stress-induced sustained increase in activity leads to loss of structural connectivity and subsequent temporal and spatial coding impairments.

## INTRODUCTION

Stress exposure impairs brain structure and function, resulting in cognitive deficits and increased risk for psychiatric disorders such as depression, schizophrenia, anxiety and post-traumatic stress disorder^1,2^. The cellular mechanisms by which stress concurs to the pathophysiology of these psychiatric disorders are not completely understood. However, it is clear that chronic stress affects the structure and physiology of the hippocampus^3–6^ and leads to long-lasting spatial memory impairments^7,8^. In rodents, structural changes include shrinking of dendritic trees and changes in density of dendritic spines and inhibitory synapses^9–19^. Physiological impairments include decreased Long Term Potentiation and increased Long Term Depression^9–13^ as well as dysregulated place cells firing in the CA1 region^14–17^. Still, the link between structural and physiological changes in the hippocampus upon stress exposure is largely unexplored, owing mostly to technical limitations.

One of these technical limitations is represented by the necessity of sacrificing the animals to study their neuronal structure, which precludes longitudinal studies during chronic stress exposure and studies of long-term structural synaptic dynamics. These dynamics have been shown to support learning^18–21^, to play an important role in the action of antidepressants^22^ and to mediate the cognitive effects of stress exposure^23^, in the neocortex. Another limitation relates to the methods used to record neuronal activity, which have confined studies investigating changes in hippocampal activity upon stress exposure to a relatively small number of place cells. This has prevented a broad understanding of the effects of repeated stress exposure on hippocampal coding. Finally, most studies focused on structure or function and utilized multiple stress paradigms and model systems to dissect structural plasticity and activity separately, thus correlating changes between connectivity and activity remained difficult.

To solve these issues, we employed deep-brain optical imaging of pyramidal neurons (PNs) located in the dorsal aspect of hippocampal CA1 (dCA1)^24^ of mice undergoing repeated stress exposure. Optical imaging enabled us to study large population of neurons at varying temporal - from hundreds of milliseconds to weeks - and spatial - from micrometers to millimeters - scales. In addition, optical imaging allowed us to perform longitudinal studies on the same subjects under different experimental conditions, thus allowing the normalization of control and experimental periods within the same individuals. Within-subjects comparison is particularly important in the hippocampus where structural connectivity and spatial coding turn over significantly^25–29^, thus it is important to take into account also changes in structure or activity patterns due to passing of time.

We studied the structural dynamics of dendritic spines - as proxies for excitatory synapses - by employing two-photon (2P) and in parallel investigated the activity patterns of dCA1 PNs - by using Wide Field Head-Mounted (WFHM) optical imaging. Repeated stress exposure led to decrease in synaptic structural connectivity which occurred in two steps: an initial decrease in spinogenesis followed by increased spine loss. Repeated stress exposure also affected patterns of activity by immediately and sustainedly increasing the amount of activity but decreasing its organization. These changes led to a significant impairment in temporal and spatial coding of dCA1 PNs.

## RESULTS

### Longitudinal tracking of dendritic spines in dCA1 PNs

By using 2P optical imaging of dCA1 in mice expressing cytoplasmic GFP in a random subset of PNs (Thy1-GFP, **Fig. 1a, b**), we longitudinally tracked basal dendrites and dendritic spines (**Fig. 1c**). After one week of baseline imaging, mice underwent 2 h of multimodal stress exposure^30^ at least 2 h prior to imaging, every day for 7 days (Repeated stress group **Fig. 1d**). These repeated stress conditions led to significant increase in the levels of plasma corticosterone (**Supplementary Fig. 1a**) and to impairments in both acquisition and recall of the Morris Water Maze (**Supplementary Fig. 1b - f**). As a control we imaged a second group of Thy1-GFP mice that underwent a single exposure to multimodal stress followed by 6 days of recovery (Acute stress group, **Fig. 1d**).

**Figure 1.**
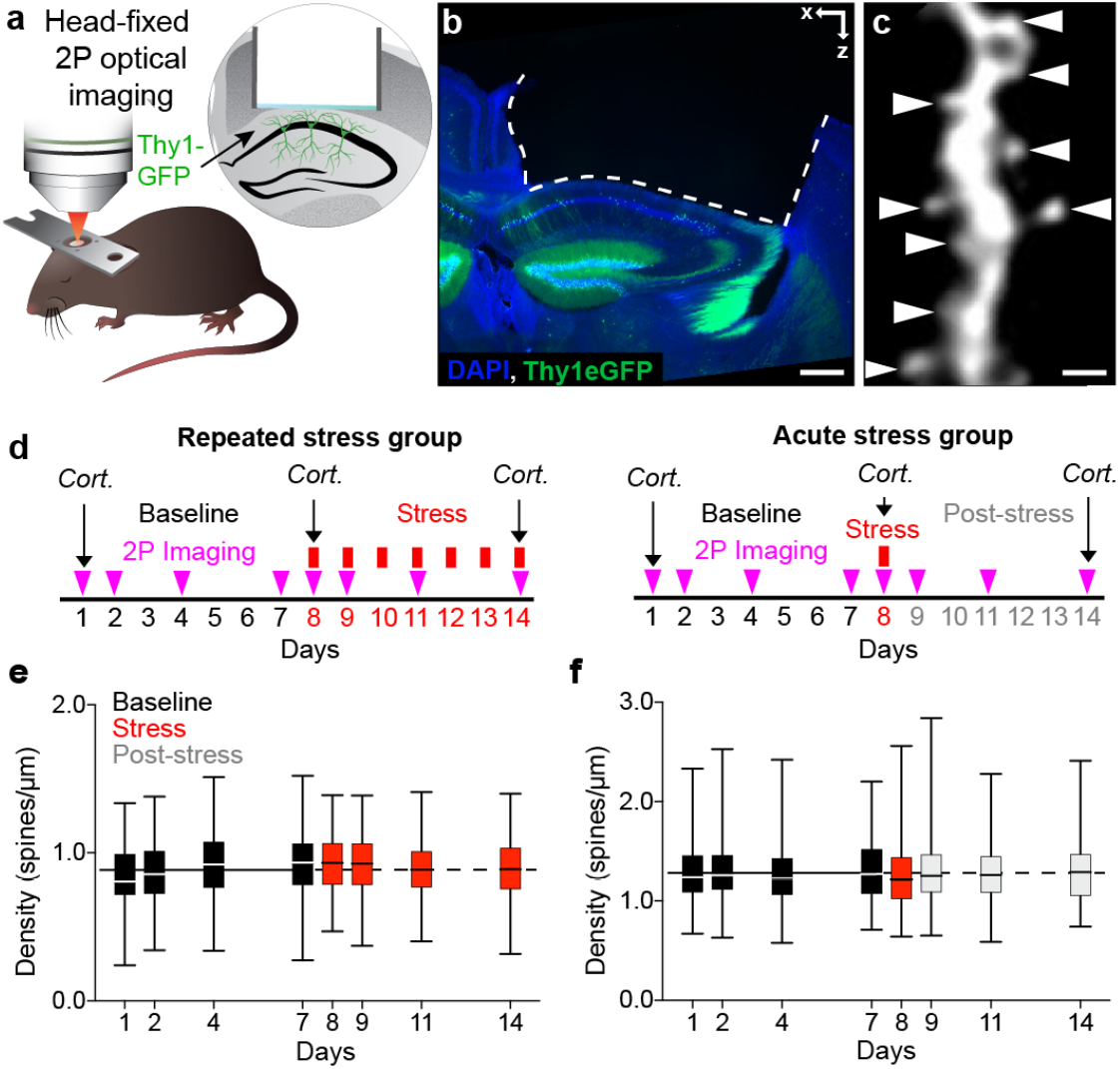
Longitudinal tracking of dendritic spines in dCA1 PNs during stress exposure. (**a**) The preparation for longitudinal 2P optical imaging of basal dendritic spines of PNs in the dCA1 of a head-fixed anesthetized mouse. (**b**) Confocal image of the dorsal hippocampus of an experimental Thy1-GFP mouse. Blue, DAPI; green, GFP. Scale bar, 200μm. (**c**) 2P image of a basal dendritic segment of a dCA1 PN of an experimental Thy1-GFP mouse (maximum intensity projection of 3 Z focal planes) showing dendritic spines (white triangles). Scale bar, 1μm. (**d**) Experimental timelines for mice stressed repeatedly (left) or acutely (right). Cort. indicates measurements of plasma Corticosterone levels. (**e** and **f**) Repeated (e) and acute (f) stress exposure did not affect spine densities. (e) p_8, 9,11, 14_ > 0.22, n_B_ = 248, n_8, 9, 11, 14_ = 114; (f) p_8, 9, 11, 14_, > 0.82; n_B_ = 352, n_8, 9, 11, 14_ = 88. Kruskal-Wallis tests against pooled baseline distribution, p values adjusted after Dunn’s multiple comparisons. Box plots: medians and quartiles of spine densities distributions per dendrite. Black solid and dashed horizontal lines: mean density of spines during baseline.

In the Repeated stress group, we tracked 114 basal dendritic segments throughout the experiment and imaged an average of 3857 (± 67 s.e.m.) and 3927 (± 41 s.e.m.) spines per session during baseline and stress respectively. In the Acute stress group, we tracked 88 basal dendritic segments and imaged 3581 (± 10 s.e.m.) and 3540 (± 27 s.e.m.) spines per session during baseline and stress/recovery respectively. The density of spines was stable during the baseline period and we found no significant changes in spine density in either the Repeated and Acute stress groups (**Fig. 1e, f**).

### Repeated stress exposure leads to an initial decrease in spinogenesis followed by increased spine loss in dCA1 PNs

We then investigated addition and persistence of spines in dCA1 PNs upon stress exposure. Repeated stress led to an immediate reduction in new spine density that persisted for at least 4 days with stress exposure (**Fig. 2a**), and, consistently, to decreased spinogenesis (**Fig. 2b**). Repeated stress also increased the fraction of spines lost but only starting from day 4 of stress exposure (**Fig. 2c**). Acute stress also led to an immediate decrease in new spine density (**Fig. 2d**), but this decrease was brief and spinogenesis recovered after stress exposure cessation (**Fig. 2e**). In contrast to repeated stress exposure, acute stress had no noteworthy effect on spine loss (**Fig. 2f**). In summary, repeated stress exposure affected dendritic spine dynamics throughout its duration - with an initial decrease in spinogenesis followed by increased spine loss (**Fig. 2g**) - while acute stress exposure only affected spinogenesis acutely (**Fig. 2h**).

**Figure 2.**
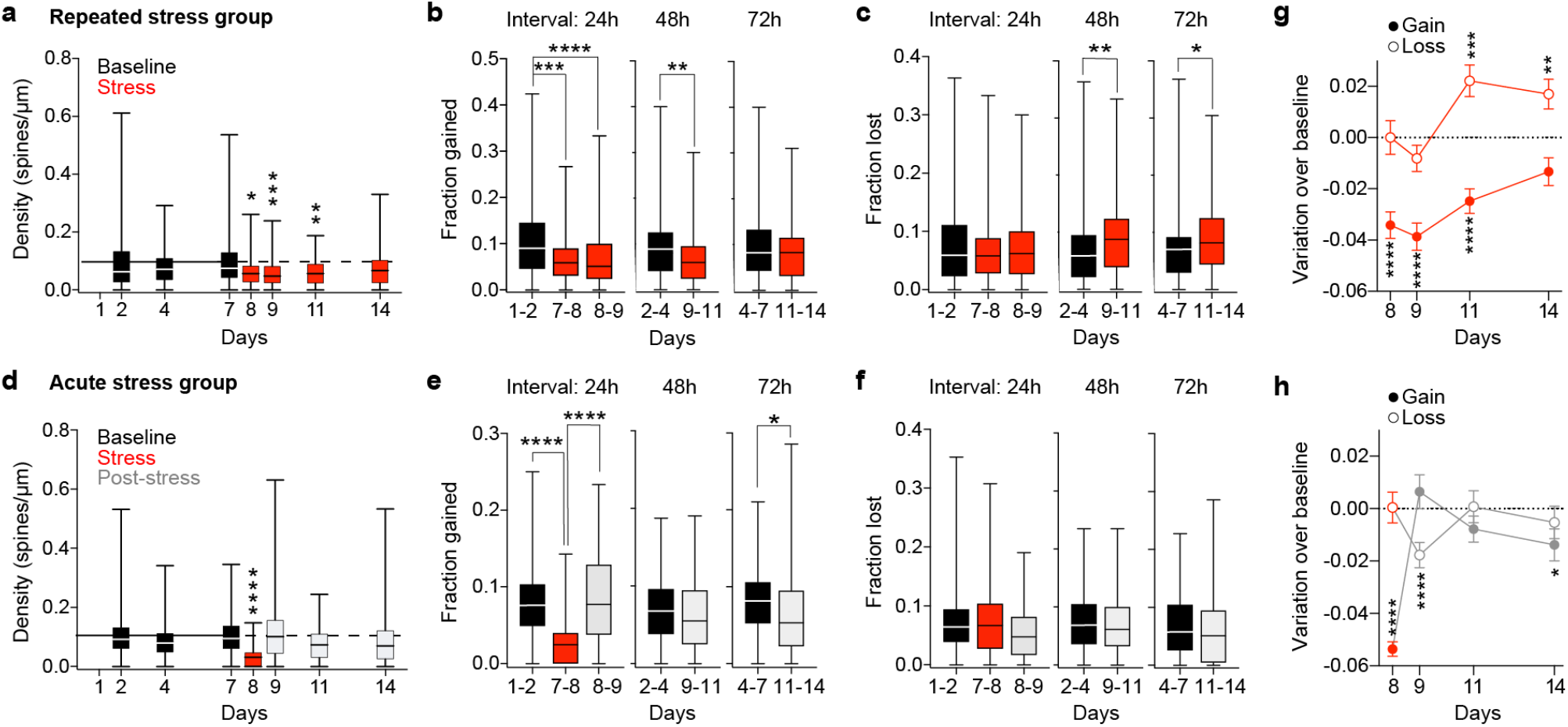
Repeated stress exposure decreases the gain and increases the loss of dendritic spines in dCA1 PNs. (**a**) The density of new spines decreased during repeated stress exposure (p_8_ = 0.025, p_9_ = 0.0002, p_11_ = 0.0018, p_14_ = 0.89, n_B_ = 248, n_8, 9, 11, 14_ = 114; Kruskal-Wallis tests against pooled baseline distribution p values adjusted after Dunn’s multiple comparisons. Black solid and dashed horizontal lines: mean density of spines during baseline. (**b**) Repeated stress exposure decreased the fraction of spines gained in the 24h and 48h intervals after the onset of the stress period (p_7-8_ =0.0001, p_8-9_ < 0.0001, p_9-11_ = 0.0022, p_11-14_ = 0.7466, n = 124; Wilcoxon matched pairs signed ranks tests to 1-2, 2-4 and 4-7 respectively, p values adjusted after Dunn’s correction for multiple comparisons). (**c**) Repeated stress exposure increased the fraction of spines lost in the 48h and 72h intervals close the end of the stress period (p_7-8_ = 0.7144, p_8-9_ > 0.9999, p_9-11_ = 0.0024, p_11-14_ = 0.0166, n = 124; Wilcoxon matched pairs signed ranks tests to 1-2, 2-4 and 4-7 respectively, p values adjusted after Dunn’s correction for multiple comparisons). Box plots: medians and quartiles of densities of new spines distributions (a) fraction of spines gained (b) or lost (c) per dendrite. (**d**) The density of new spines decreased immediately after single stress exposure but recovered after stress interruption (p_8_ < 0.0001, p_9_ > 0.999, p_11_ = 0.057, p_14_ = 0.05, n_B_ = 352, n_8, 9, 11, 14_ = 88; Kruskal-Wallis tests against pooled baseline distribution, p values adjusted after Dunn’s correction for multiple comparisons). Black solid and dashed horizontal lines: mean density of spines during baseline. (**e**) Acute stress decreased the fraction of spines gained immediately after stress exposure (p_7-8_ < 0.0001, p_8-9_ > 0.9999, p_9-11_ = 0.3483, p_11-14_ = 0.0182, n = 88; Wilcoxon matched pairs signed ranks tests to 1-2, 2-4 and 4-7 respectively, p values adjusted after Dunn’s correction for multiple comparisons). (**f**) Acute stress had no effect on spine loss (f, p_7-8_ > 0.9999, p_8-9_ = 0.3263, p_9-11_ = 0.8860, p_11-14_ = 0.2665, n = 88; Wilcoxon matched pairs signed ranks tests to 1-2, 2-4 and 4-7 respectively, p values adjusted after Dunn’s correction for multiple comparisons). Box plots: medians and quartiles of the distributions of densities of new spines (e) fraction of spines gained (f) or lost (g) per dendrite. (**g**) Repeated stress exposure decreased spine gain initially and later increased spine loss (gain, p_7-8_ < 0.0001, p_8-9_ < 0.0001, p_9-11_ < 0.0001, p_11-14_ = 0.07, n = 124; loss, p_7-8_ > 0.99, p_8-9_ = 0.11, p_9-11_ = 0.0005, p_11-14_ = 0.0043, n = 124; One sample t-test against the value 0). (**h**) Acute stress decreased spine gain immediately and later increased spine loss (gain, p_7-8_ < 0.0001, p_8-9_ = 0.33, p_9-11_ = 0.12, p_11-14_ = 0.026, n = 124; loss, p_7-8_ = 0.94, p_8-9_ = 0.0004, p_9-11_ = 0.91, p_11-14_ = 0.39, n = 124; One sample t-test against the value 0). Circles: mean variation per dendrites of spine gain (full) or loss (empty) over baseline of the corresponding time window.

### Repeated and acute stress have opposite effects on stabilization of spines

Consistent with increased spine loss, we found a decrease in spine survival (All spines, **Fig. 3a**) upon repeated stress exposure. Loss of synaptic stability affected spines that already existed before the stress event (Preexisting spines, **Fig. 3b**) and new spines detected on the first (**Fig. 3c**, left) and second (**Fig. 3c**, right) days of stress exposure. In contrast, a single exposure to stress increased the survival of spines (All spines, **Fig. 3d**). Increased synaptic stability, however, was not due to pre-existing spines (**Fig. 3e**) but to new spines detected specifically on the day of stress exposure (**Fig. 3f**).

**Figure 3.**
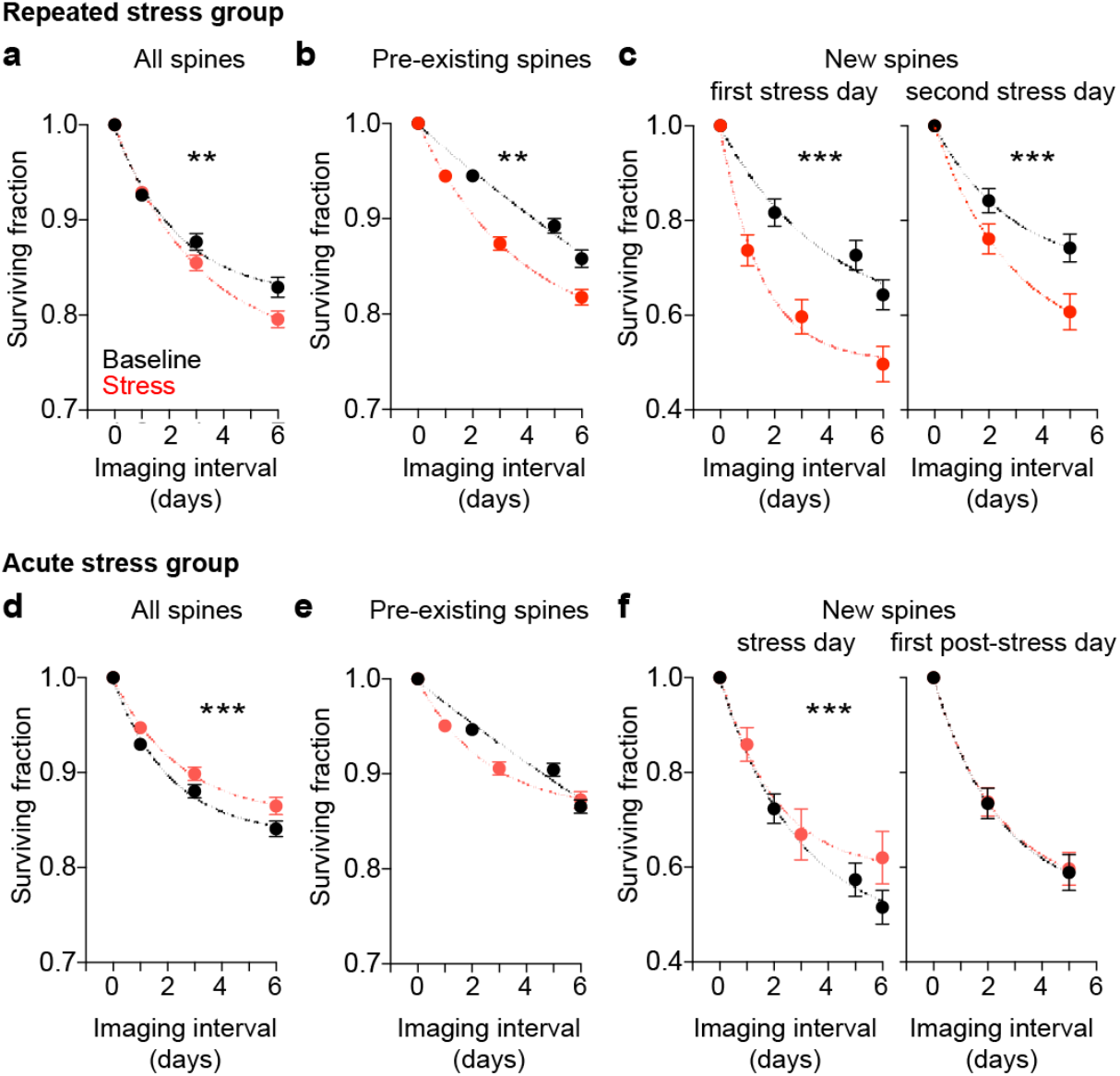
Repeated stress exposure decreases the survival of dendritic spines in dorsal CA1 PNs. (**a − c**) Repeated stress exposure decreased the survival of all (a), pre-existing (b) and new (c) spines detected on the first (left) and second (right) days of stress. p_All_ = 0.003, p_Pre-existing_ = 0.001, p_NewFirstday_ < 0.001, p_NewSecondday_ < 0.001, n_All_ = 372, n_Pre-existing_ = 372, n_NewFirstday_ = 312, n_NewSecondday_ = 130. (**d – f**) A single exposure to stress slightly but significantly increased the survival of all spines (d), but it did not affect pre-existing spines (e). Increase in spine survival was due to increased survival rate of new spines detected on the day of stress (f, left) but not on the following day (f, right). p_All_ < 0.001, p_Pre-existing_ = 0.251, p_NewFirstday_ < 0.001, p_NewSecondday_ = 0.861, n_All_ = 264, n_Pre-existing_ = 264, n_NewFirstday_ = 177, n_NewSecondday_ = 154. Mann-Whitney U test between all non-1 Baseline versus Stress points. Circles: mean surviving fractions per dendrite. Error bars: s.e.m. Curves: single exponential decays fit to the data.

Thus, while repeated stress exposure destabilizes all spines, acute stress stabilizes specifically the spines born in temporal proximity to stress.

### Imaging activity of hundreds of neurons in freely-moving mice undergoing repeated exposure to stress

We expected changes in synaptic connectivity to affect significantly patterns of neuronal activity, we thus investigated activity patterns of dCA1 PNs in a separate group of animals undergoing the same repeated stress paradigm.

By using WFHM fluorescence microscopy we studied the activity of dCA1 PNs expressing the Ca^2+^ sensor GCamP6f (**Fig. 4a**). We imaged Ca^2+^ transients in several hundred dCA1 PNs (**Fig. 4b**) in mice exploring a circular arena for 15 minutes every day for 7 days (**Fig. 4c**). We then divided the mice in two groups: the repeated stress group underwent 2 h of multimodal stress exposure at least 2 h prior to every imaging session every day for 7 days (**Fig. 4c**), while the control (No Stress) group was imaged every day for 7 additional days without stress exposure (**Fig. 4c**). We imaged up to 1356 neurons in a single session and an average of 393 (± 24 s.e.m.), 468 (± 78 s.e.m.) and 405 (± 50 s.e.m.) neurons per mouse per session during the baseline, control and stress periods respectively. The number of neurons did not significantly change over time or across groups (**Supplementary Fig. 2a, b**).

**Figure 4.**
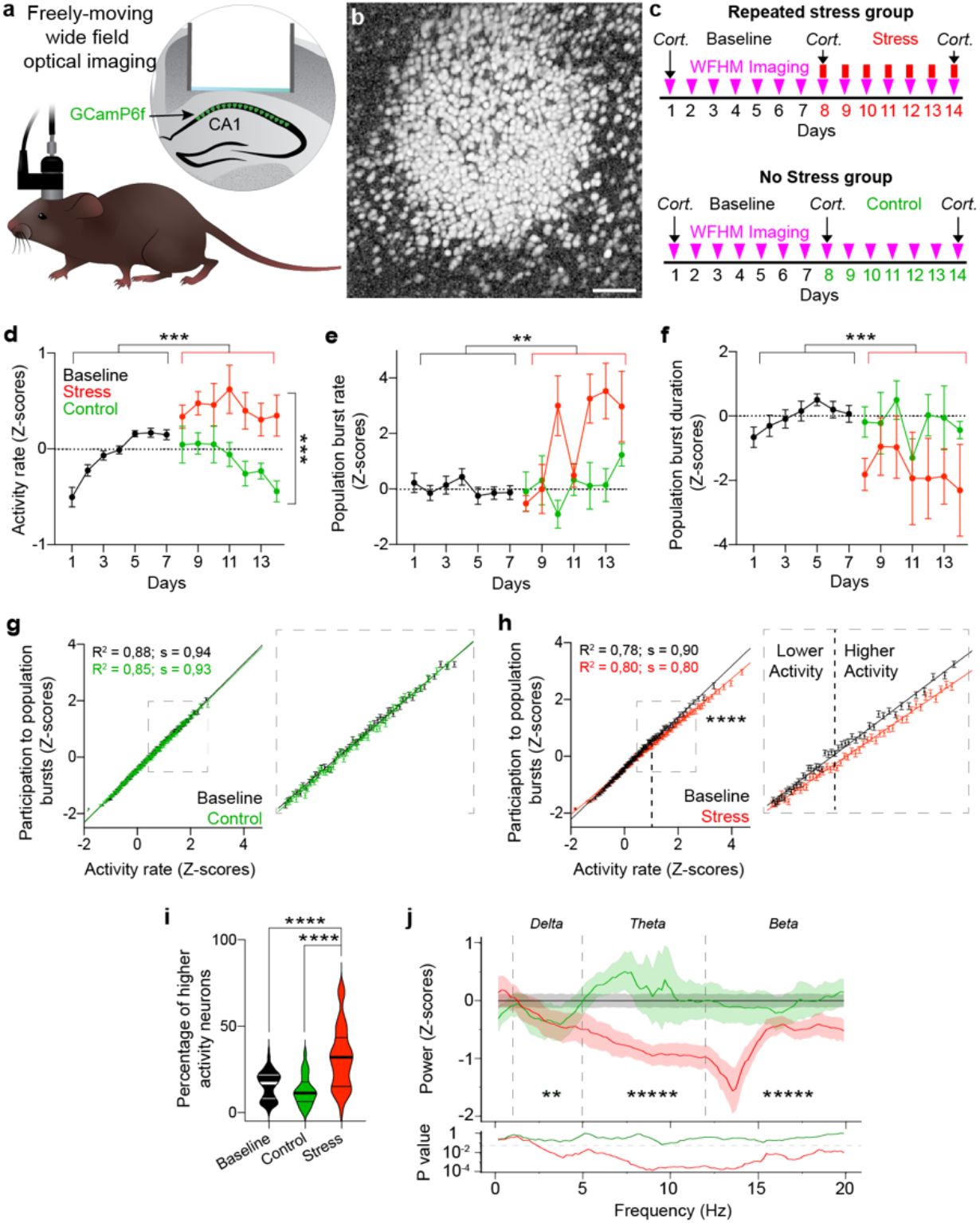
Exposure to repeated stress induces hyperactivity but reduces the theta coordination in dCA1. (**a**) The preparation for repeated WFHM optical imaging of neuronal activity in the dCA1 of freely-moving mice. (**b**) Neurons active during a 15 minute-long exploration of a round arena. The image shows the pixel-wise correlation map across one recording session. Scale bar, 80 μm. (**c**) Experimental timelines for mice stressed repeatedly (top) or not stressed (bottom). Cort. indicates measurements of plasma corticosterone levels. (**d – f**) Stress exposure increased neuronal activity rates (d) and incidence rate of hippocampal population bursts (e), but it decreased the duration of the population bursts (f). d: p_B-C_ = 0.2, p_B-S_ < 0.0001, p_C-S_ < 0.001; n_B_ = 86, n_C_ = 27, n_S_ = 49. e: p_B-C_ > 0.99, p_B-S_ = 0.004, p_C-S_ = 0.19; n_B_ = 89, n_C_ = 27, n_S_ = 49. f: p_B-C_ > 0.99, p_B-S_ = 0.0012, p_C-S_ = 0.19; n_B_ = 86, n_C_ = 27, n_S_ = 50. Kruskal-Wallis test, p values adjusted after Dunn’s corrections for multiple comparisons. Circles: mean activity rate (d), population burst rate (e) and population burst duration (f) per mouse, Z-scored over each mouse. Error bars: s.e.m. (**g** and **h**) Activity rate correlated with participation in population bursts of PNs (g: R^2^_B_ = 0.88, R^2^_C_ = 0.85; p_B_ < 0.0001, p_C_ < 0.0001; n_B_ = 13862, n_C_ = 13057. h: R^2^_B_ = 0.78, R^2^_S_ = 0.80; p_B_ < 0.0001, p_S_ < 0.0001; n_B_ = 20664, n_S_ = 19858). Stress exposure changed this relationship (p_B-S_ < 0.00001, n_B_ = 13862, n_S_ = 19858; 2-way ANOVA) due to a reduction in participation of neurons with higher activity rates (neurons with activity rate > 1 Z-score, g and h insets). Circles: participation index of neurons in population bursts binned over activity rates. Lines: linear fit to the data. Error bars: s.e.m. (**i**) Stress exposure doubled the percentage of neurons with higher activity rates (p_B-C_ = 0.4, p_B-S_ < 0.0001, p_C-S_ < 0.0001; n_B_ = 87, n_C_ = 28, n_S_ = 49; Kruskal-Wallis test, p values adjusted after Dunn’s corrections for multiple comparisons). Violin plots: median and quartiles of the percentage of neurons with higher activity rates distributions. (**j**) Stress exposure decreased neuronal synchronization in the Delta (1-5Hz), Theta (5-12 Hz) and Alpha (1220Hz) bands (Delta: p_B-C_ < 0.00001, p_B-S_ < 0.00001, p_C-S_ = 0.0094; n_B_ = 1357, n_C_ = 529 n_S_ = 506. Theta: p_B-C_ = 0.0006, p_B-S_ < 0.00001, p_C-S_< 0.00001; n_B_ = 2400, n_C_ = 920, n_S_ = 960. Alpha: p_B-C_ < 0.00001, p_B-S_ < 0.00001, p_C-S_ < 0.00001; n_B_ = 2700, n_C_ = 1035, n_S_ = 1080; Mann-Whitney U test). Asterisks: p values for the comparisons of Control and Stress. Bottom pane: p values of the comparisons of Control (green) and Stress (red) powers with Baseline (bin width = 0.17 Hz) throughout the spectrum.

Repeated stress exposure increased the levels of circulating corticosterone - without, however, reaching statistical significance over baseline (**Supplementary Fig. 3a – c**) - and affected exploration behavior by decreasing average speed, increasing immobility and decreasing occupancy of the arena (**Supplementary Fig. 3d – l**). Stress exposure did not affect permanence in the center of the arena (**Supplementary Fig. 3m – o**).

### Exposure to repeated stress induces hyperactivity and reduces the temporal coordination in dCA1

Exposure to repeated stress increased PNs’ activity rate (**Fig. 4d**) and the rate of population bursts (**Fig. 4e**) but it decreased the duration of population bursts (**Fig. 4f**). Repeated stress reduced the participation of neurons with higher activity rate to population bursts (**Fig. 4g and h**). Setting the point at which the baseline and stress linear relationships started to differ (activity rate > 1 Z-score) as a threshold for higher activity rate PNs (**Fig. 4h** inset), we found that repeated stress exposure almost doubled the number of PNs with higher activity rates (**Fig. 4i**). To investigate synchronization of dCA1 PNs, we calculated the power spectra of the population activity using Fast Fourier Transform and normalized them to the average spectrum of the baseline. Repeated stress exposure significantly decreased neuronal synchronization in the Delta (1-5Hz), Theta (5-12 Hz) and Beta (12-20Hz) frequency bands, with the largest difference between control and stress groups being in the Theta band (average control 0.2 Z-score above baseline, average stress 0.8 Z-score below baseline) (**Fig. 4j**).

These data show that while repeated stress exposure increases the amount of activity in hippocampal dCA1, it impairs the temporal coordination of this activity.

### Repeated stress exposure decreases pairwise temporal correlations and leads to co-activation networks with more modular and assortative topology

To further study the effects of stress exposure on the temporal coordination of dCA1 PNs’ activity, we calculated the pairwise temporal correlation among neurons. While the correlation increased during the second experimental week in the control group, stress exposure prevented this increase (**Fig. 5a**) both for PNs with lower and higher activity rates (**Fig. 5b**). The difference between control and stress groups became significant after 5 days of repeated stress both for PNs with lower and higher activity rate (**Fig. 5c**).

**Figure 5.**
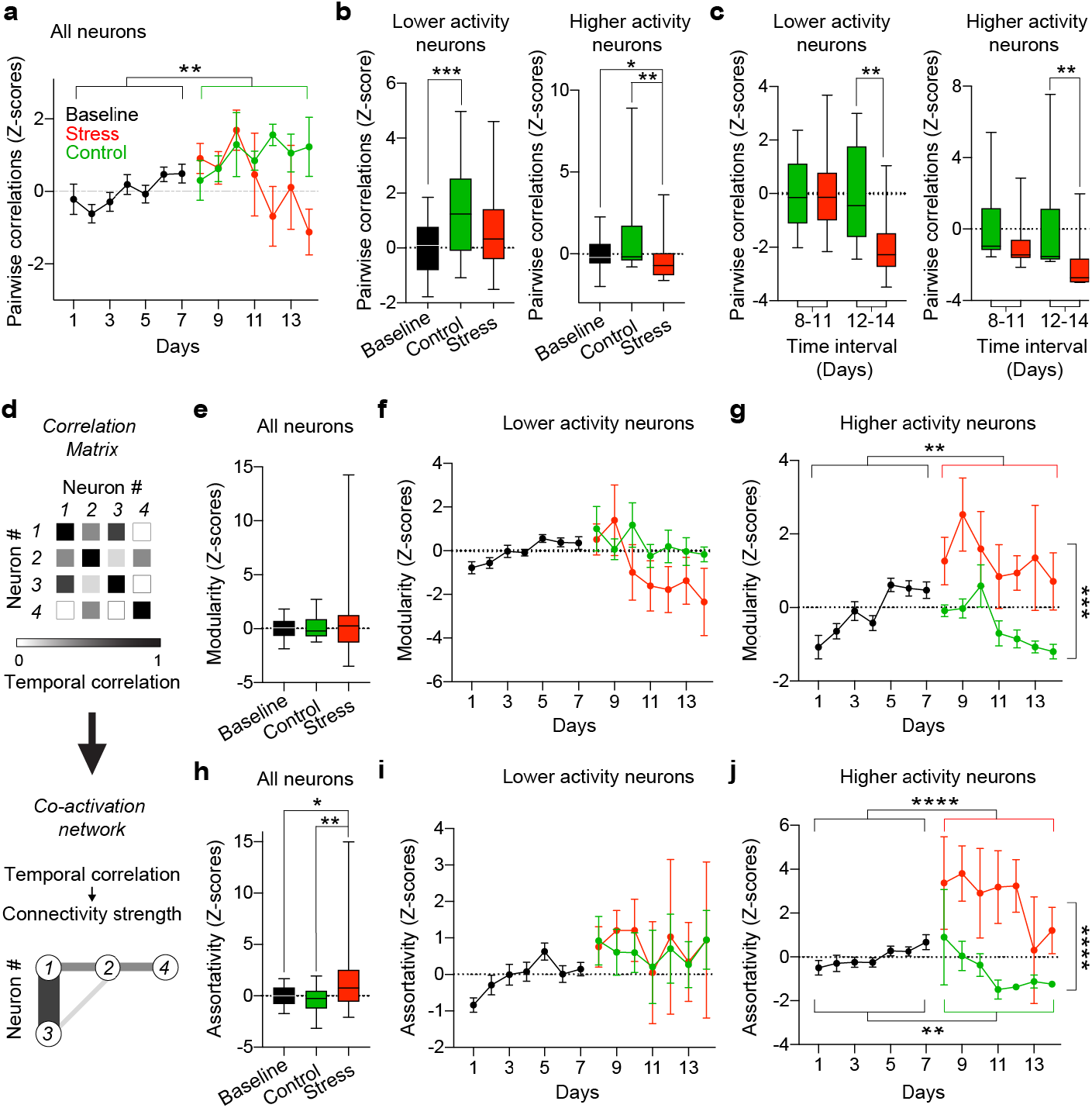
Exposure to repeated stress decreases pairwise temporal correlations and leads to co-activation networks with more modular and assortative topology. (**a**) Stress exposure blocked increase in pairwise temporal correlations in all neurons (p_B-C_ = 0.002, p_B-S_ = 0.25, p_C-S_ = 0.21; n_B_ = 75, n_C_ = 25, n_S_ = 42). (**b**) Correlation of lower activity neurons significantly increased during the second week of imaging in the control group (left: p_B-C_ = 0.0008, p_B-S_ = 0.083, p_C-S_ = 0.3; n_B_ = 75, n_C_ = 25, n_S_ = 42). The difference between Control and Stress groups became significant after 5 days of repeated stress (right: p_8-11_ >0.99, p_12-14_ = 0.0038; n_C 8-11_ = 16, n_S 8-11_ = 28, n_C 12-14_ = 9, n_S 12-14_ = 14). (**c**) Stress exposure decreased the temporal correlation of higher activity neurons (left: p_B-C_ = 0.36, p_B-S_ = 0.025, p_C-S_ = 0.002; n_B_ = 75, n_C_ = 24, n_S_ = 42). The difference between Control and Stress groups became highly significant after 5 days of repeated stress (right: p_8-11_ = 0.11, p_12-14_ = 0.0039; n_C 8-11_ = 15, n_S 8-11_ = 28, n_C 12-14_ = 9, n_S 12-14_ = 14). Data in each time interval were normalized to the mean of the control group in that interval. (**d**) Schematic description of the transformation from correlation matrices to co-activity networks. (**e - g**) Stress exposure did not affect modularity of networks comprising all neurons (e) and lower activity rate neurons (f) but significantly increased modularity of the network comprising only higher activity neurons (g). d: p_B-C_ > 0.99, p_B-S_ > 0.99; p_C-S_ > 0.99 n_B_ = 86, n_C_ = 28, n_S_ = 49. e: p_B-C_ = 0.99, p_B-S_ = 0.12, p_C-S_ = 0.20; n_B_ = 86, n_C_ = 28, n_S_ = 49. f: p_B-C_ = 0.14, p_B-S_ = 0.009, p_C-S_ = 0.0001; n_B_ = 79, n_C_ = 25, n_S_ = 47. (**h - j**) Stress exposure increased assortativity of the network comprising all neurons (h) mostly due to increased assortativity of the network comprising only higher activity neurons (I, j). g: p_B-C_ = 0.65, p_B-S_ = 0.022, p_C-S_ = 0.005; n_B_ = 86, n_C_ = 85, n_S_ = 49. h: p_B-C_ = 0.18, p_B-S_ = 0.62, p_C-S_ > 0.99; n_B_ = 86, n_C_ = 28, n_S_ = 49. i: p_B-C_ = 0.006, p_B-S_ < 0.0001, p_C-S_ < 0.0001; n_B_ = 79, n_C_ = 25, n_S_ = 47. Circles: mean pairwise correlations (a), modularity (f, g) or assortativity (i, j) per mouse, Z-scored over each mouse. Error bars: s.e.m. (**b** and **c**) Box plots: medians and quartiles of the distributions of mean pairwise correlation (b and c) modularity (e) or assortativity (h) per mouse per session, Z-scored over each mouse. Kruskal-Wallis test, p values adjusted after Dunn’s corrections for multiple comparisons.

We then generated co-activation networks whose nodes were the PNs and whose edges - or connectivity strengths - were the values of pairwise temporal correlation between neurons (**Fig. 5d**). Repeated stress exposure increased the propensity of higher activity rate neurons to cluster into modules (Modularity, **Fig. 5e-g**) and the propensity of neurons to interact with neurons of similar connectivity degree (Assortativity, **Fig. 5h**) mostly due to neurons with higher activity rates (**Fig. 5i, j**).

These data show that repeated stress exposure decreases the temporal correlation of dCA1 PNs’ activity but increases co-activity of neurons with higher activity rates leading to relative decoupling of higher and lower activity neurons in co-activation networks.

### Repeated stress exposure increases the number of ensembles in dCA1 but it lowers effective size and reactivation potential of these ensembles

We then used principal component analysis to extract activity patterns corresponding to neural ensembles and defined significant ensembles as ensembles whose encoding strength during a session was ≥ 1^31^ (**Fig. 6a**). Repeated stress exposure increased the fraction of significant ensembles comprising only higher activity rate neurons (**Fig. 6b**). For all significant ensembles, we sorted neurons according to their weights and used the width of the weight’s distribution at half its maximum as a threshold to demarcate the ensembles’ cores from their peripheries (**Fig. 6c**). Repeated stress exposure decreased the size of ensembles’ cores, mainly due to the contribution of higher activity rate neurons (**Fig. 6d**). While participation of neurons to ensembles’ cores increased in control groups (**Fig. 6e**), stress exposure prevented this increase (**Fig. 6e**), mostly by decreasing the mean participation of higher activity rate neurons to ensembles’ cores (**Fig. 6f**) and increasing the fraction of neurons in ensembles’ peripheries (**Fig. 6g**). The ensemble’s encoding strength also increased in the control groups, but stress exposure also prevented this increase (**Fig. 6h**). Moreover, neurons participating to ensembles with encoding strengths similar to control showed activity rates higher than control upon stress exposure (**Fig. 6i**).

**Figure 6.**
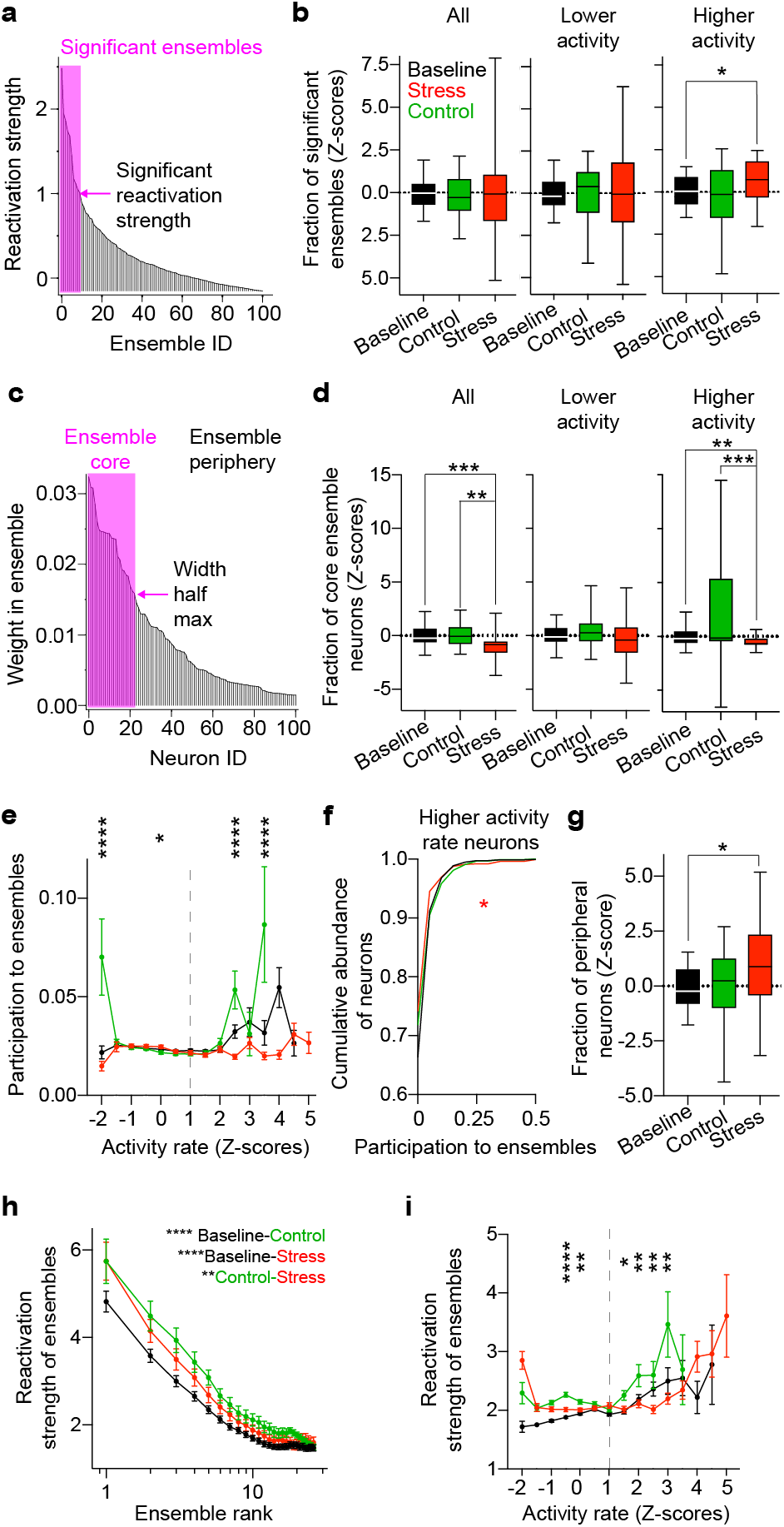
Exposure to repeated stress increases the number of ensembles in dCA1 but lowers effective size and reactivation potential of these ensembles. (**a**) Neuronal ensembles in one mouse during an exploration session sorted according to their normalized reactivation strengths. The arrow denotes the threshold for reactivation strength = 1, the pink rectangle indicates the significant ensembles corresponding to reactivation strength > 1. (**b**) Stress exposure did not change the fraction of significant ensembles among all (left) and lower activity neurons (middle) but significantly increased the fraction of significant ensembles among higher activity neurons (right). All: p_B-C_ > 0.99, p_B-S_ > 0.99, p_C-S_ > 0.99; n_B_ = 84, n_C_ = 28, n_S_ = 45. Lower activity: p_B-C_ > 0.99, p_B-S_> 0.99, p_C-S_ > 0.99; n_B_ = 83, n_C_ = 28, n_S_ = 43. Higher activity: p_B-C_ > 0.99, p_B-S_ = 0.03, p_C-S_ = 0.17; n_B_ = 50, n_C_ = 15, n_S_ = 32. (**c**) Neurons in one mouse during an exploration session, sorted according to their weights within an ensemble. The arrow indicates the threshold that demarcates the ensemble’s core (pink rectangle). (**d**) Stress exposure decreased the size of ensembles’ cores for all neurons (left: p_B-C_ > 0.99, p_B-S_ < 0.0001, p_C-S_ = 0.0017; n_B_ = 87, n_C_ = 24, n_S_ = 38), mainly due to higher activity neurons. Lower activity: p_B-C_ = 0.84, p_B-S_ = 0.38 p_C-S_ = 0.09; n_B_ = 83, n_C_ = 28, n_S_ = 40. Higher activity: p_B-C_ = 0.25, p_B-S_ = 0.0002, p_C-S_ = 0.09; n_B_ = 80, n_C_ = 26, n_S_ = 39. Box plots: medians and quartiles of mean fraction of significant ensembles (b) or of neurons participating in ensembles’ cores (d) distributions per mouse per session. Kruskal-Wallis test, p values adjusted after Dunn’s corrections for multiple comparisons. (**e**) Participation of neurons in ensembles’ cores increased with time, but stress exposure prevented this increase (p_−2_ < 0.0001, p_−1.5, −1, −0.5_ > 0.28, p_0_ = 0.015, p_0.5, 1, 1.5, 2_ > 0.38, p_2.5_ < 0.0001, p_3_ = 0.78, p_3.5_ < 0.0001; n_−2_ > 68, n_−1.5, −1, −0.5_ > 702, n_0_ > 2461, p_0.5, 1, 1.5, 2_ > 285, p_2.5_ > 75, p_3_ > 18, p_3.5_ > 5; Pairwise, two-tailed t-tests between control and stress per each activity rate bin. Circles: mean ensemble participation index binned over activity rates. Error bars: s.e.m. Vertical dashed line indicates the threshold for neurons with higher activity rates. (**f**) Stress exposure decreased the mean participation of higher activity rate neurons to ensembles’ cores (p_B-C_ > 0.99, p_B-S_ = 0.031, p_C-S_ = 0.32; n_B_ = 4445, n_C_ = 1201, n_S_ = 5863; Kruskal-Wallis test, p values adjusted after Dunn’s corrections for multiple comparisons). (**g**) Stress exposure increased the fraction of neurons in ensembles’ peripheries (p_B-C_ > 0.99, p_B-S_ = 0.046, p_C-S_ = 0.28; n_B_ = 40, n_C_ = 28, n_S_ = 35; Kruskal-Wallis test, p values adjusted after Dunn’s corrections for multiple comparisons). Box plots: medians and quartiles of mean fraction of neurons participating in ensemble’s periphery distributions per mouse per session. (**h**) Reactivation strength of ensembles increased with time, but stress exposure prevented this increase (p_B-C_ < 0.0001, p_B-S_ < 0.0001, p_C-S_ = 0.0072; n_B_ = 1462, n_C_ = 510, n_S_ = 749; Two-Way ANOVA). Circles: mean reactivation strengths of ensembles index ranked according to decreasing values. Error bars: s.e.m. (**i**) Stress exposure increased the activity rates of neurons participating in ensembles with similar reactivation strengths (p_−2, −1.5, −1_ > 0.06, p_−0.5_ < 0.00001, p_0_ = 0.004, p_0.5, 1_ > 0.22, p_1.5_ = 0.013, p_2_ = 0.003, p_2.5_ = 0.006, p_3_ = 0.006, p_3.5_ = 0.61; n_−2, −1.5, −1_ > 30, n_−0.5_ > 1036, n_0_ > 1141, n_0.5, 1_ > 873, n_1.5_ > 354, n_2_ > 136, n_2.5_ > 42, n_3_ > 8, n_3.5_ > 4; Pairwise, two-tailed t-tests between control and stress per each activity rate bin. Circles: mean reactivation strengths of the ensembles in which each neuron took part binned over activity rates. Error bars: s.e.m. Vertical dashed line indicates the threshold for neurons with higher activity rates

In summary, these results reveal that repeated stress exposure leads to smaller neuronal ensembles with lower recurrence power.

### Repeated stress exposure decreases the amount of spatial information and impairs spatial coding in dCA1

To investigate the effects of repeated stress on spatial coding, we calculated the amount of Spatial Information^32^ (SI) per neuron in a total of 25.230, 9.575 and 11.668 neurons during baseline, control and stress periods respectively. Repeated stress exposure decreased the amount of SI (**Fig. 7a**). During baseline lower activity rates were associated with higher SI and higher activity rates were associated with lower SI. In control animals this sigmoidal relationship showed a trend towards increase of SI in lower activity rate neurons and decrease of SI in higher activity rate neurons. Repeated stress exposure, however, decreased SI in lower activity rate neurons and increased SI in higher activity rate neurons (**Fig. 7b**). Stress exposure affected the relationship between SI and the participation to population bursts in a similar fashion, by decreasing the participation of neurons with lower SI and increasing the participation of neurons with higher SI (**Fig. 7c**). Thus, repeated stress exposure impaired the relationship between activity rates, SI and participation in population bursts (**Fig. 7d**).

**Figure 7.**
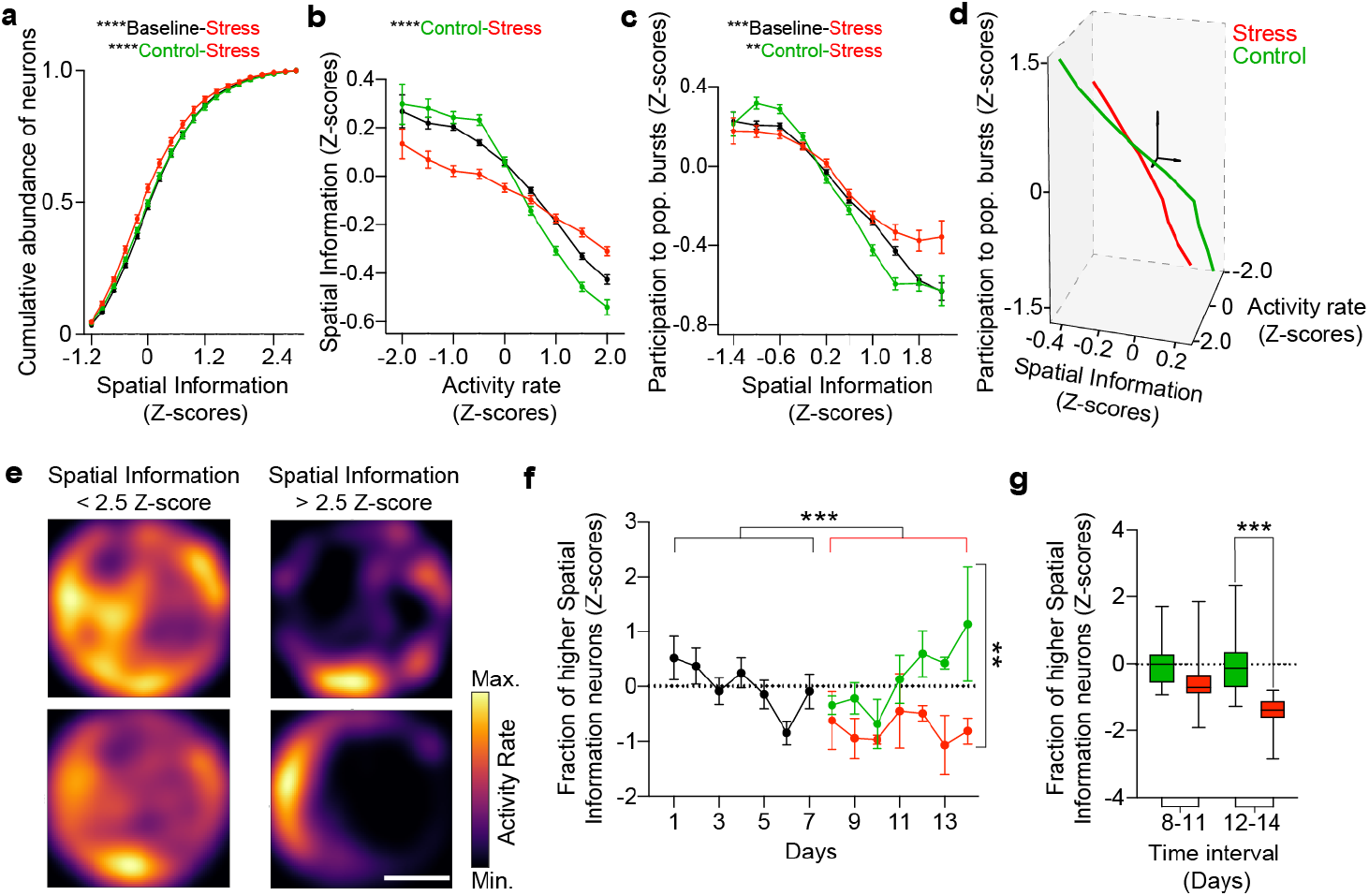
Repeated stress exposure decreases the amount of spatial information in dCA1. (**a**) Stress exposure decreased the amount of SI per neuron (p_B-C_ = 0.38, p_B-S_ < 0.0001, p_C-S_ < 0.0001; n_B_ = 1152, n_C_ = 396, n_S_ = 540; 2-way ANOVA, p values adjusted after Tukey’s corrections for multiple comparisons). (**b**) Stress exposure affected the relationship between rate of activity and SI by decreasing the SI in neurons with lower activity rate and increasing the SI in neurons with higher activity rate (p_B-C_ = 0.21, p_B-S_ = 0.074, p_C-S_ < 0.0001; n_B_ = 9575, n_C_ = 25230, n_S_ = 1168; 2-way ANOVA, p values adjusted after Tukey’s corrections for multiple comparisons). circles: mean SI binned over activity rates. Error bars: s.e.m. (**c**) Stress exposure affected the relationship between SI and the rate of participation in ensembles by decreasing the rate of participation in significant ensembles in neurons with lower SI and increasing the rate of participation in significant ensembles in neurons with higher SI (p_B-C_ = 0.49, p_B-S_ = 0.0005, p_C-S_ < 0.0022; n_B_ = 9575, n_C_ = 25230, n_S_ = 1168; 2-way ANOVA, p values adjusted after Tukey’s corrections for multiple comparisons). circles: mean ensemble participation index binned over activity rates. Error bars: s.e.m. (**d**) Stress exposure shifted the state of the dCA1 PNs network in the three-dimensional space defined by activity, SI and participation in ensembles. The arrows in the center represent the axes originating at 0 with the arrows pointing in the positive direction. (**e**) Example firing maps of two neurons with lower SI (left) and two neurons with higher SI (right) recorded in the same mouse during the same session. Scale bar, 20 cm. (**f**) Stress exposure decreased the fraction of neurons with higher spatial information (p_B-C_ > 0.99, p_B-S_ = 0.0006, p_C-S_ = 0.0011; n_B_ = 59, n_C_ = 26, n_S_ = 29; Kruskal-Wallis test, p values adjusted after Dunn’s corrections for multiple comparisons). Circles: mean fraction of higher SI neurons per mouse per session, Z-scored over each mouse baseline period. Error bars: s.e.m. (**g**) The difference between control and stress groups became significant after 5 days of repeated stress (p_8-11_ = 0.075, p_12-14_ = 0.0007; n_C 8-11_ = 16, n_S 8-11_ = 20, n_C 12-14_ = 11, n_S 12-14_ = 9; Kruskal-Wallis test, p values adjusted after Dunn’s corrections for multiple comparisons). Box plots: medians and quartiles of the mean fractions of higher SI neuron distributions per mouse per session, Z-scored over each mouse. Data in each time interval were normalized to the mean of the control group in that interval.

We then investigated the effect of repeated stress exposure on place cell-like activity by focusing on neurons with the highest SI (SI ≥ 2.5 Z-score, **Fig. 7e**). Stress exposure decreased the fraction of high-SI neurons (**Fig. 7f**) with the difference between control and stress groups becoming highly significant only after 5 days of repeated stress (**Fig. 7g**).

Overall, repeated stress exposure impairs spatial coding in dCA1 by disrupting the relationship between activity rates, SI and participation in population bursts and preventing the increase in place cell-like activity.

## DISCUSSION

We employed 2P deep-brain time lapse and WFHM optical imaging to track synaptic connectivity and activity patterns of dCA1 PN’s respectively, during a baseline period and repeated or acute stress exposure longitudinally in the same individuals. We detected multiple effects of stress on structural dynamics and activity patterns (**Fig. 8a**).

**Figure 8.**
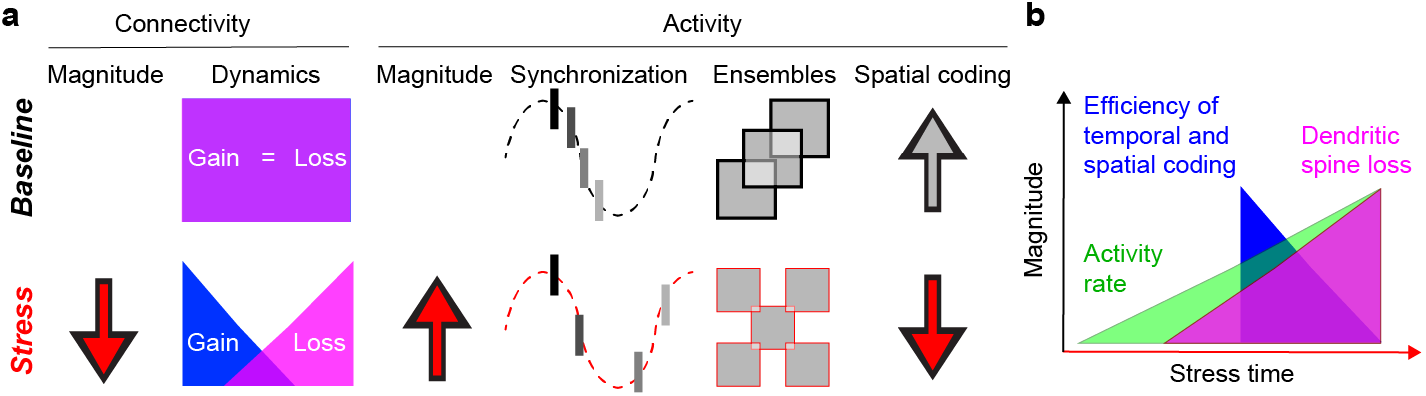
Graphical summary of the results. (**a**) Stress exposure affected connectivity and activity patterns of dCA1 PNs. (**b**) Temporal sequence of stress-induced impairments in connectivity and activity patterns of dCA1 PNs.

### Decreased spinogenesis precedes increased spine loss upon exposure to repeated stress

Despite average stability of spine density, we found two different effects on synaptic dynamics. First, spinogenesis decreased after single stress exposure and lasted for up to 4 days of repeated stress exposure (**Fig. 2b and e**). This might depend on non-genomic effects of glucocorticoids and corticotropin releasing factor^4^ and is consistent with the fact that stress decreases Long-Term Potentiation (LTP)^9,11,33^ and LTP induction promotes spinogenesis^34,35^. Decreased spinogenesis, however, was transient and recovered to baseline levels after seven days of repeated stress exposure, suggesting the presence of compensatory mechanisms under sustained stress conditions. Second, spine loss became apparent after 4 days of repeated but not after single stress exposure (**Fig. 2c**) and, concomitantly, activity rate steadily increased during repeated stress exposure (**Fig. 5d**). These results suggest that a sustained increase in activity is necessary for significant spine loss, possibly due to cytotoxic effects or homeostatic compensation of hyperactivity^36^.

Previous work showed a small but significant decrease in dCA1 spine density upon stress exposure^37,38^. However, in our hands the density of spines was stable during the baseline period and no significant changes occurred after repeated or single exposure to stress. This discrepancy might be due to intrinsic differences between the different animal models used (mice *vs*. rats) or to the dCA1 layer analyzed (basal *vs*. apical).

### Changes in structural synaptic connectivity and impaired activity patterns upon repeated stress exposure

Most excitatory presynaptic terminals in the basal aspect of dCA1 PNs originate in CA2 and CA3^39^, thus loss of excitatory synapses on dCA1 PNs implies compromised information transfer from CA2 and CA3 and possibly decreased feedforward inhibition on dCA1 PNs. In fact, loss of excitatory connectivity on dCA1 PNs might also parallel loss of excitatory connectivity on neighboring inhibitory neurons. Altogether, this might lead to decreased or disorganized activity of dCA1 inhibitory neurons as previously shown^23,40^. Further studies will need to dissect the effects of stress on the connectivity of specific pre-and post-synaptic partners. However, given the role of Parvalbumin-expressing (PV+) inhibitory neurons in synchronizing CA1 PNs^41,42^, decreased or impaired activity of PV+ inhibitory neurons could lead to lower synchronization in dCA1 PNs specifically in theta range^43,44^. This is consistent with the decrease in synchrony of CA1 activity - which was more pronounced in the theta range (**Fig. 4j**) - we found upon repeated stress exposure and suggests that decrease in structural excitatory connectivity on dCA1 PNs leads to decreased synchrony in activity through disruption of PV+ inhibitory neuronal activity.

Moreover, increased spine loss preceded decreased correlation of activity and number of neurons with higher SI (**Fig. 8b**) but became apparent only after several days of hyperactivity of dCA1 PNs upon repeated stress exposure. Although we cannot exclude that these changes occur independently, the temporal sequence of these phenomena (**Fig. 8b**) suggests that stress-induced sustained increase in activity leads to loss of structural connectivity leading in turn to impairments in temporal and spatial coding.

Altogether our findings suggest a role in structural plasticity in mediating the effects of repeated stress on dCA1 activity patterns and coding.

### Stress-induced stabilization of new spines in dCA1 might support learning of stress-related events

Acute and repeated stress exposure showed opposite effects on stability of excitatory synaptic connectivity. While repeated stress exposure decreased overall spine survival (**Fig. 3a - c**), a single exposure to stress significantly increased the survival of new spines born in temporal vicinity to stress exposure (**Fig. 3d**). Such stabilization could depend on stress-specific regulation of adhesion molecules such as NCAM or L1^45^ and also on increased trafficking of AMPA receptors due to non-genomic action of glucocorticoids^46,47^. By analogy to the neocortex, where stabilization of new dendritic spines can support acquisition of new memories^18–21^, we suggest that the increase in stability of new spines born in temporal proximity to stress exposure might be a cellular mechanism supporting learning of information related to the stressful event^48^. Sustained stress exposure, however, - possibly due to cytotoxic or homeostatic effects - impairs spine survival thus pushing the system towards impaired learning^48^.

### The number of highly active neurons increases with stress exposure but their activity is disorganized

Repeated stress exposure led increased activity in dCA1 PNs (**Fig. 8a**), in line with previous reports^23,30,49^. However, the temporal organization of this activity differed from control, in that neurons with higher activity rates participated less in bursts, segregated into modules but were excluded from the cores of ensembles. Highly active neurons are a smaller fraction of all active neurons^50^, that tends to be stably active through time^51^, can be preferentially recruited into engrams^52,53^ and possibly dominate information transfer in the hippocampus and other cortical regions^50,54^. Thus, despite their smaller abundance, disruption of the temporal organization of highly active neurons might have larger effects on hippocampal representations and potentially underlie the stress-induced impairment in learning and memory.

### Repeated stress exposure impairs the temporal correlation and the temporal coding of dCA1 PNs

Temporal correlation or synchronization of neuronal activity is thought to support information transfer across different brain areas and within groups of neurons in the same brain area as well as learning. In the hippocampus, synchronization in the theta frequency has been implicated in local information processing^32,55–57^, long-range communication to neocortical areas^58,59^ and synaptic plasticity^60^. In our experiments, repeated exploration of the same environment led to increased theta power and pairwise correlation of neuronal activity consistent with previous work^58,61^. Repeated exposure to stress, however, lowered synchronization and pairwise correlations and prevented the increase in theta power (**Fig. 8a**).

Pyramidal neurons in CA1 can show sequential activity within a few hundred milliseconds giving rise to neuronal ensembles^32,62,63^. Ensemble activity is a form of temporal organization thought to underpin information representation^64,65^ and memory formation^66–69^. In addition, the strength of reactivation of these neuronal ensembles is linked to their importance in hippocampal learning and recall^69,70^. Repeated stress exposure decreased the size of ensembles and prevented increase in the ensemble’s encoding strength (**Fig. 8a**). Disruption in temporal correlation and coding of dCA1 PNs might underlie the stress-induced deficits in hippocampal dependent learning and memory.

### Repeated stress exposure impairs spatial coding of dCA1 PNs

Previous studies investigated the effect of stress exposure on spatial coding in rodents focused on the activity of place cells^13,15–17^. Here, we opted for a more inclusive approach by investigating the amount of SI per neuron in hundreds of neurons per animal. Stress exposure not only reduced the mean amount of SI but it also significantly impaired the efficiency of spatial coding. In fact, while repeated exploration of the arena led to more SI being encoded with lower activity rates - a more energy-efficient way to encode space - stress led to less spatial information being coded with higher activity rates - a less efficient spatial coding (**Fig. 8a**).

In summary, our data suggest a causal relationship between increase in neuronal activity, spine loss and impaired temporal and spatial coding in the dorsal hippocampal CA1 of mice, possibly due to disorganized activity of local inhibitory neurons. However, further work will be necessary to support this causal relationship. Specifically, it will be important to find ways to manipulate synaptic stability to either rescue changes in temporal or spatial coding due to synaptic destabilization during stress exposure or mimic the effects of stress on in temporal or spatial coding through synaptic destabilization in absence of stress-induced hyperactivity.

## ACKNOWLEDGEMENTS

Alon Chen is supported by an FP7 Grant from the ERC, the ERANET and I-CORE programs, the Israeli Ministry of Health, the BMBF, the Max Planck Foundation, the Nella and Leon Benoziyo Center for Neurological Diseases, the Henry Chanoch Krenter Institute for Biomedical Imaging and Genomics, The ISF, the Perlman Family, the Adelis, Marc Besen, Pratt and Irving I. Moskowitz foundations and by Roberto and Renata Ruhman, Bruno and Simone Lich. Alessio Attardo is supported by the Max Planck Society, the Deutsche Forschung Gemeinschaft (DFG) and Schram Foundation.

## ONLINE METHODS

### Subjects

All animals used were either C57BL6 (for longitudinal WFHM freely-moving optical imaging) or Thy1-GFP transgenic (for longitudinal 2P head-fixed optical imaging) male mice between 3 and 6 months of age. Animals had free access to food and water with a 12/12 light/dark cycle. All animal procedures conformed to the guidelines of the Max Planck Society and the Animal authority, Regierung Oberbayern (Regierung von Oberbayern – Veterinärwesen) and were approved in the Tierversuchslizenz - ROB-55.2Vet-2532.Vet_02-16-48.

### Viral vector injection

Intracranial injections of Adeno Associated Viral suspensions were carried out according to standard methods^32^. We injected 500 nL of a viral suspension (AAV2/1.Syn.GCaMP6f.WPRE.SV40; titer, ~10^12^ genomes copy/ml; U Penn Vector Core) at a rate of 100 nL/min at two separate locations in dCA1 (AP, −2 mm and −2.5 mm; ML, 1.6 mm and 2.1 mm relative to bregma; DV, 1.3 mm). Mice were allowed to recover for a minimum of 10 days.

### Implantation of a chronic hippocampal imaging cannula

We implanted chronic hippocampal cannulas according as previously described^28^. An imaging cannula (2.5 mm-inner-diameter, 1.6 mm-long) was inserted into a 3 mm-diameter craniotomy (AP, −2.3 mm; ML, 2 mm relative to bregma) applying slight pressure onto the tissue to stabilize the preparation. The cannula was fixed and sealed to the skull using Metabond (Parkell). A custom head plate was positioned and fixed with dental cement.

*For longitudinal 2P head-fixed optical imaging*, the head plate consisted of a thin aluminum or titanium slab with holes to fit the imaging cannula and a holder positioned on the 2P microscope stage.

*For longitudinal WFHM freely-moving optical imaging*, the head plate consisted of a plastic cylinder threaded on the inside surface to enable mounting of the microscope objective and the microscope itself (Doriclenses).

### In vivo optical imaging

*For longitudinal 2P head-fixed optical imaging*, mice were anesthetized with 2.5% Isoflurane and placed under the microscope (Bruker Ultima IV) onto a 37°C heating pad (CMA 450) while the head was fixed via a head plate holder. During imaging mice were kept under constant anesthesia (1.5% Isoflurane). We aligned the imaging cannula to the light path as previously described^***29***^. To image eGFP-expressing dendrites and dendritic spines we used a 25x water immersion objective (Olympus XLPlan N 25x/1.00 SVMP) and a pulsed infrared laser tuned to 920 nm. We acquired z-stacks (48.18 μm^2^ single section area, 1 μm z-step, 5-60 z-steps, zoom 10x, 28.6-115.5 mW laser power at the sample) of the dendrite of interest using a resonant scanner at each time point.

*For longitudinal WFHM freely-moving optical imaging*, mice were familiarized with the recording room and the circular arena for 3 days. During this period the objective lens (0.5 NA, 2.4 WD, Air immersion) and the microscopes (Doriclenses, S model) were mounted on the head of the animals, and the animals were free to explore the arena for 5-10 minutes / day. On the last day of habituation we rotated the objective lens to reach a depth at which we could detect neuronal activity in the raw images. We then glued the objective in place using epoxy glue and removed the miniaturized microscope before placing the animals back in their home cage. During the 14 days following the habituation period, we mounted the miniaturized microscope to the imaging objective, placed the animals in the same arena each day and recorded Ca^2+^ transients as proxy for neuronal activity for 15 minutes using 488 nm continuous wavelength laser (Thorlabs) illumination (1mW average power) at a sampling rate of 45 Hz using commercial software (Doric Neuroscience Studio software). We cleaned the arena and changed the bedding every day and mixed oat flakes in the bedding material to motivate exploration.

### Pre- and post-processing, data reduction and definition of metrics used for analysis

*For longitudinal 2P head fixed optical imaging*, we compensated for motion artifacts and scored as previously described^29^. In **Fig. 2**, fractional gain and fractional loss between two consecutive imaging points were defined as the number of spines gained or lost between the first and the second time points, normalized by the number of spines present in the first time point. In **Fig. 3**, fractional survival was defined as the number of spines surviving between the first and each of the other time points, normalized by the number of spines present in the first Baseline time point, first and second stress timepoints.

*For longitudinal WFHM freely-moving optical imaging*, we first performed data reduction using CaImAn ^24^ We then spatially down-sampled the image time series by a factor of 2 and motion corrected using the NoRMCorre algorithm^71^. We simultaneously denoised, deconvolved, and demixed our data using the extended version of the constrained non-negative matrix factorization (CNMF-E) algorithm^72^. We finally time-stamped the deconvolved signal for all neurons. The parameters used for the CNMFE algorithm are reported in **Supplementary table 1**. In **Fig. 4**, *activity rates* of neuronal populations were calculated as the sum of individual neuronal activity rate per time bin (22 ms). *Population bursts* were defined as events in which the population rate was 1 standard deviation (SD) above the mean and peaks at least to 2.5 SD above the mean. *Population burst rate* was the number of population bursts recorded in each session divided by the total recording time. The *population burst duration* was the time span where population rate was above mean + 1SD threshold. *Participation to population bursts* was defined as the fraction of population bursts that a neuron contributes to, normalized to the total number of population bursts. The *power spectral density* of population activity rates was obtained by the Fast Fourier Transform algorithm implemented in Spectrum software package^73^. In **Fig. 5**, we calculated the pairwise Pearson correlation matrices per each mouse at each time point by using neuronal activity time series binarized and binned within 132ms bins. We then generated network adjacency matrices by binarizing the correlation matrices in comparison to 95 percentile correlation values for correlations among shuffled data set of neurons with scrambled activity rate. Network *Modularity and Assortativity* were calculated using Newman’s spectral community detection^74^ implemented in Brain Connectivity Toolbox^75^. In **Fig. 6**, we defined *neuronal ensembles* as the principal components of the pairwise Pearson correlation matrices^76^ and their *encoding strengths* as the ratio between ensemble eigenvalue (*λ*) and the Marcenko-Pastur upper threshold *λ_max_*.

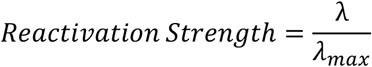

Significant ensembles were defined as ensembles with eigenvalues exceeding Marcenko-Pastur threshold^31^; only significant ensembles were considered for further analyses. We defined the *ensemble core* as the fraction of neurons with weights greater than half of maximum weight for a given ensemble. We defined *the participation index to ensembles* of a neuron as the number of significant ensembles having that neuron as a part their core divided by total number of significant ensembles in the recording session. In **Fig. 7**, to obtain *spatial activity maps* we determined the location of the animal at the onset of each calcium event and we divided the arena into 5 cm x 5 cm non-overlapping bins. We then calculated the activity rate in each bin as the number of calcium events for each bin divided by the occupancy time of that bin. Only calcium events where animal speed exceeded 2cm/s were considered in the analysis. *Mean activity rates* were calculated by averaging the activity rate of each neuron. *Spatial information* was calculated as previously described^31^

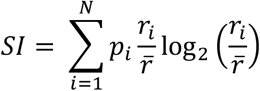

Where *p_i_* is the occupancy probability, *r_i_* is the average activity rate of the i-th bin and 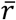 is the average firing rate of the neuron. The spatial information value was Z-scored to the distribution of spatial information values obtained by shuffling the activity times of the neuron 100 times.

### Multimodal stress exposure

Multimodal stress exposure was performed as previously described^32^. Briefly mice were placed into an open plastic tube to restrain their movement and allow breathing. Cage mates were placed on a rocking platform with light and exposed to light and loud noise at the same time for a total duration of two hours, once a day for a total of seven days.

### Analysis of animal behavior

Morris Water Maze. The animal behavior was recorded during navigation of a standard (circular, filled with milky water at a temperature of 24 degrees Celsius) Water Maze arena. The position and speed of animals was extracted from video recording feed using Anymaze (Stoelting). Latency and quadrant occupancy were calculated using Anymaze (Stoelting), while navigation strategies were extracted using custom Matlab software^30^.

Exploration of the open arena. The animal behavior was recorded simultaneously with neuronal recording during 15 minutes-long exploration of a circular arena (40 cm diameter). The position and speed of animals was extracted from a live video feed using a custom made Bonsai^77^ script. Animal’s trajectories were corrected manually for transient tracking errors.

### Statistical analysis

We used Pearson’s correlations, Kruskal-Wallis tests with Dunn’s corrections for multiple comparisons, 2-way ANOVA with Tukey’s corrections for multiple comparisons, Mann-Whitney U tests, Wilcoxon matched pairs signed ranks tests with Dunn’s correction for multiple comparisons, one sample t-tests. Statistical analysis and plotting were done with Prism 8 (GraphPad) or Python software.* p ≤ 0.05, ** p ≤ 0.01, *** p ≤ 0.001, **** p < 0.0001.

### Data and software availability

The analysis code can be found on Github. All original raw data will be made available by the corresponding authors upon request.

