## Supplementary Figures and Captions for "Aberrant structural synaptic dynamics precede disorganization of temporal and spatial coding in the hippocampus upon repeated stress exposure"

[4] Current address: Queensland Brain Institute, The University of Queensland, Brisbane, QLD 4072, Australia.

[*] Equal contribution

### SUPPLEMENTARY FIGURES AND CAPTIONS

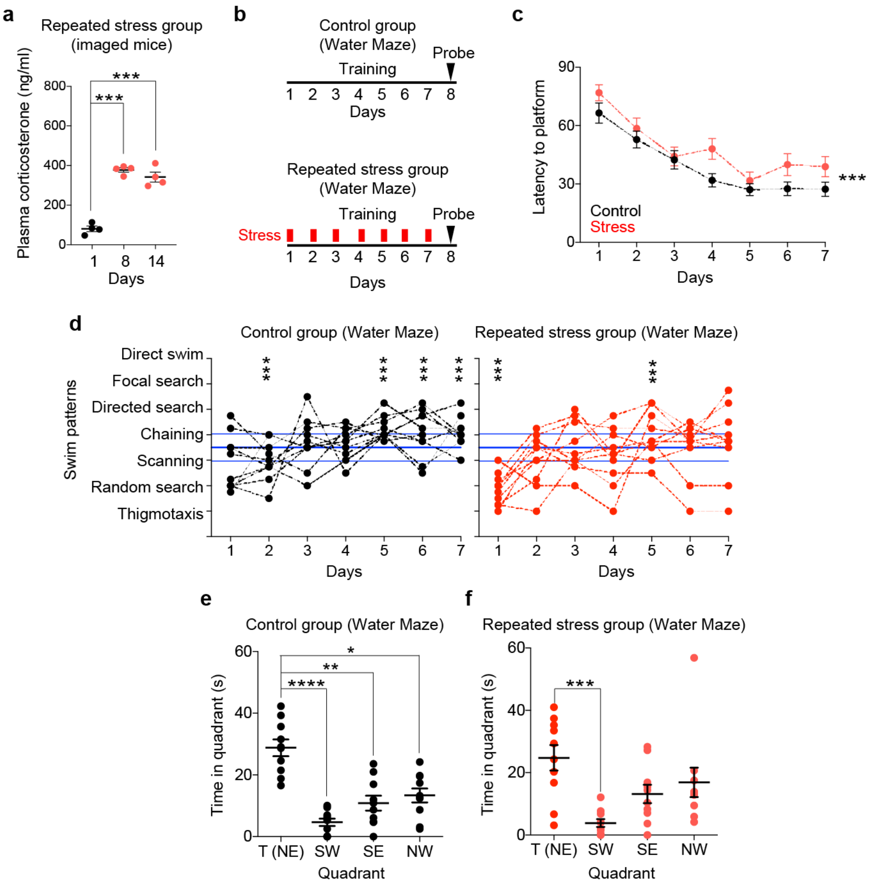


**Supplementary Figure 1 | Repeated stress increases the blood corticosterone levels and impairs learning and recall of an hippocampal dependent memory task.**

**(a)** Repeated stress increased the levels of corticosterone in the blood of mice undergoing longitudinal in vivo imaging ($p_{1\text{-}8} = 0.0001$, $p_{1\text{-}14} = 0.0007$, n = 12; Repeated Measurements ANOVA, p values adjusted after Sidak's corrections for multiple comparisons).

(**b**) Timeline of Morris Water maze training and testing.

(**c**) Repeated stress during training increased latency to reach the hidden platform ($p < 0.0001$, n = 80; Repeated Measurements ANOVA). Circles: average latency of 10 mice per time point. Black: control group, red: stress group.

(**d**) Left: control mice showed progression towards swim strategies indicative of learning and at the end of the training period were on average significantly above criterion ($p_{1,3,4} > 0.9999$; $p_{2,5,6,7} < 0.0001$; binomial test on rank sum p-value; n = 10 animals per day; Binomial test on rank sum p-value). Right: mice stressed in each training day showed a slower progression towards swim strategies indicative of learning and at the end of the training period were not on average significantly above criterion ($p_{2, 3, 4, 6, 7}$ >0.9999; $p_{1, 5}$ < 0.0001; binomial test on rank sum p-value; n = 10 animals per day; Binomial test on rank sum p-value). Circles: average strategy of 4 trials per mouse. Black: control group, red: stress group. Blue lines: average strategy per day (solid) with s.e.m. (dashed) assuming equal occurrence of each strategy each day.

(**e**) Control mice spent significantly more time in the target quadrant (NE), during the probe trial ($p_{SW}$<0.0001, $p_{SE}$=0.004, $p_{NW}$=0.036; n = 10 animals per quadrant; Kruskal-Wallis test, p values adjusted after Dunn's corrections for multiple comparisons).

(**f**) Mice stressed repeatedly during training spent significantly less time only in the quadrant opposite to the target quadrant, during the probe trial ($p_{SW}$=0 .002, $p_{SE}$=0 .29, $p_{NW}$= 0.68; n = 10 animals per quadrant; Kruskal-Wallis test, p values adjusted after Dunn's corrections for multiple comparisons).

Circles: time per quadrant per mouse. Black: control group, red: stress group. Horizontal lines: means ± s.e.m.

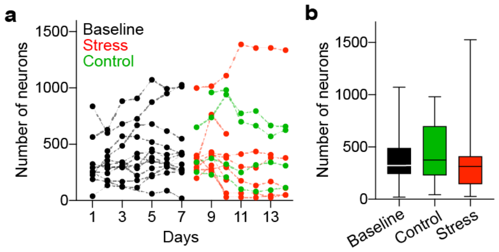


**Supplementary Figure 2 | The number of neurons recorded by WFHM optical imaging is stable through conditions.**

(**a, b)** The number of neurons imaged by WFHM microscopy did not change through baseline, control and stress periods ($p_{B\text{-}C} = 0.83$, $p_{B\text{-}S} > 0.9999$, $p_{C\text{-}S} = 0.27$; $n_B = 106$, $n_C = 35$, $n_S = 60$; Kruskal-Wallis test, p values adjusted after Dunn's corrections for multiple comparisons).

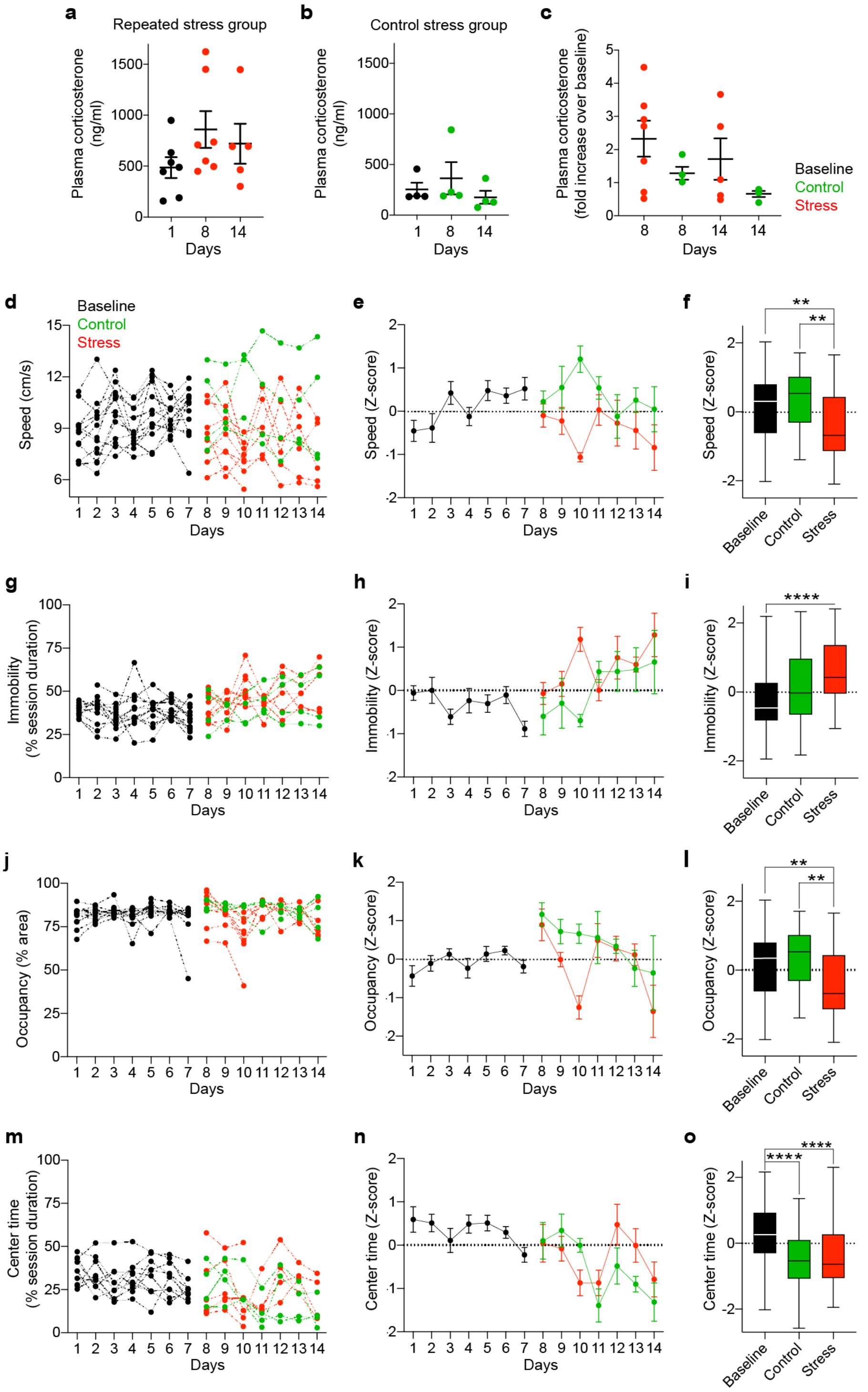


**Supplementary Figure 3 | Repeated stress concomitantly to WFHM optical imaging does not significantly increase the blood corticosterone levels but impairs several aspects of free navigation in a circular arena.** (**a**) Repeated stress did not significantly increase the levels of corticosterone in the blood of mice undergoing WFHM optical imaging ($p_{1\text{-}8} = 0.12$, $p_{1\text{-}14} = 0.70$, $n_1 = 7$, $n_8 = 7$, $n_{14} = 5$; Kruskal-Wallis test, p values adjusted after Dunn's corrections for multiple comparisons).

(**b**) The levels of corticosterone in the blood of mice undergoing WFHM optical imaging did not significantly increase in the control group ($p_{1-8} = 0.31$, $p_{1-14} = 0.31$, $n_1 = 4$, $n_8 = 4$, $n_{14} = 4$; Kruskal-Wallis test, p values adjusted after Dunn's corrections for multiple comparisons).

(**c**) Stressed animals showed a fold increase in the levels of corticosterone in their blood that did not reach significance over controls ($p_{S-C,8} > 0.9999$, $p_{S-C,14} = 0.43$; $n_8 = 11$, $n_{14} = 9$; Kruskal-Wallis test, p values adjusted after Dunn's corrections for multiple comparisons).

(**d - f**) Repeated stress decreased average speed during free exploration of the arena ($p_{B-C} = 0.77$, $p_{B-S} = 0.0048$; $p_{C-S} = 0.0023$; $n_B = 90$, $n_C = 27$, $n_B = 90$; $n_S = 50$; Kruskal-Wallis test, p values adjusted after Dunn's corrections for multiple comparisons).

(**g - i**) Repeated stress increased immobility during free exploration of the arena ($p_{B-C} = 0.20$, $p_{B-S} < 0.0001$; $p_{C-S} = 0.22$; $n_B = 90$, $n_C = 27$, $n_B = 90$; $n_S = 50$; Kruskal-Wallis test, p values adjusted after Dunn's corrections for multiple comparisons).

(**j - l**) Repeated stress decreased average occupancy of the arena ($p_{B-C} = 0.77$, $p_{B-S} = 0.0048$; $p_{C-S} = 0.0023$; $n_B = 90$, $n_C = 27$, $n_B = 90$; $n_S = 50$; Kruskal-Wallis test, p values adjusted after Dunn's corrections for multiple comparisons).

(**m - o**) Repeated stress did not affect permanence in the center of the arena ($p_{B-C} = 0.001$, $p_{B-S} < 0.0001$; $p_{C-S} > 0.999$; $n_B = 90$, $n_C = 27$, $n_B = 90$; $n_S = 43$; Kruskal-Wallis test, p values adjusted after Dunn's corrections for multiple comparisons).
